# Midbody remnant inheritance is regulated by the ESCRT subunit CHMP4C

**DOI:** 10.1101/810523

**Authors:** Javier Casares-Arias, María Ujué Gonzalez, Alvaro San Paulo, Leandro N. Ventimiglia, Jessica B. A. Sadler, David G. Miguez, Leticia Labat-de-Hoz, Armando Rubio-Ramos, Laura Rangel, Miguel Bernabé-Rubio, Jaime Fernández-Barrera, Isabel Correas, Juan Martín-Serrano, Miguel A. Alonso

## Abstract

The inheritance of the midbody remnant (MBR) breaks the symmetry of the two daughter cells, with functional consequences for lumen and primary cilium formation by polarized epithelial cells, and also for development and differentiation. However, despite their importance, neither the relationship between the plasma membrane and the inherited MBR nor the mechanism of MBR inheritance is well known. Here, the analysis by correlative light and ultra-high-resolution scanning electron microscopy reveals a membranous stalk that physically connects the MBR to the apical membrane of epithelial cells. The stalk, which derives from the uncleaved side of the midbody, concentrates the ESCRT machinery. The ESCRT CHMP4C subunit enables MBR inheritance, and its depletion dramatically reduces the percentage of ciliated cells. We demonstrate: (1) that MBRs are physically connected to the plasma membrane, (2) how CHMP4C helps maintain the integrity of the connection, and (3) the functional importance of the connection.

## Introduction

The midbody (MB) is the narrow bridge that connects the two nascent daughter cells resulting from animal cell division. MB cleavage results in the physical separation of the cells, through a process known as abscission, and in the formation of an MB remnant (MBR) (Fededa and Gerlich, 2012, Mierzwa and Gerlich, 2014). Increasing evidence indicates that, instead of being an abscission byproduct, the MBR assumes important roles in development and differentiation (Kuo et al., 2011, Singh and Pohl, 2014, Ettinger et al., 2011, Peterman et al., 2019). In polarized renal epithelial cells, the MBR licenses the centrosome to assemble the primary cilium, which is a solitary plasma membrane protrusion involved in the regulation of multiple developmental signaling pathways (Bernabe-Rubio et al., 2016), and defines the location of the apical membrane during lumen formation (Lujan et al., 2017).

The MB is continuous with the plasma membrane and consists of an electron-dense central region called the Flemming body (FB) (Byers and Abramson, 1968), which comprises anti-parallel microtubule bundles. Flanking the FB, the MB has two arms, containing parallel microtubule bundles, vesicles and protein factors, that bridge the two daughter cells. In principle, when severing occurs on both arms, the MBR becomes extracellular and it can remain free in the extracellular milieu, or stay attached to the surface of one of the daughter cells or of a neighboring cell, or be eliminated. However, severing on just one arm should lead to the MBR to be inherited by the cell on the opposite side, although this has not been well documented experimentally (Schiel et al., 2011). Given the importance of the MBR (Peterman and Prekeris, 2019, Chen et al., 2013), it seems inevitable that its fate must be tightly regulated (Ou et al., 2014, Dionne et al., 2015). However, despite the enormous effort expended on trying to understand the mechanism of the first cleavage of the MB, which marks the end of cell division, little attention has been paid to the inheritance of the connected MBRs and, thus, to the regulation of the cut of the membrane of the other MB arm.

In this study, using ultra-high-resolution scanning electron microscopy (SEM), we demonstrate the existence of physical continuity between the MBR membrane and the plasma membrane of Madin-Darby canine kidney (MDCK) cells, and show that only one side of the MB is cleaved in most cases. We find that, once abscission is completed, the charged multivesicular body protein (CHMP) 4C subunit of the endosomal sorting complex required for transport (ESCRT) complex delays the cleavage of the membrane of the other arm, allowing the MBR to remain on the cell surface as an organelle physically connected to the rest of the plasma membrane. The connection enables the MBR to license the centrosome for primary cilium assembly, and might be also important in other processes involving the MBR.

## Results

### MBRs of MDCK cells are connected to the plasma membrane by a membranous extension

Epithelial MDCK cells constitute a paradigm of polarized epithelial cell (Rodriguez-Boulan et al., 2005). Given the important role of the MBR in MDCK cells, we chose this cell line as a model cell system to study whether there was continuity between the MBR and the rest of the cell. Unlike tumor-derived cell lines (Kuo et al., 2011, Ettinger et al., 2011), MDCK cells have a single MBR at most (Bernabe-Rubio et al., 2016). Quantitative analysis indicates that >95% of MBRs are on the apical surface (Figures 1A and S1A). Mitotic kinesin-like protein 1 (MKLP1), which localizes to the FB, is a key component of the centraspindlin complex, responsible for antiparallel microtubule bundling during anaphase, MB stability and ESCRT recruitment during cytokinesis (Mishima, 2016, Zhao et al., 2006). Super-resolution structured illumination microscopy showed that MBRs are formed by the FB, which was visualized by staining endogenous MKLP1, flanked by two small microtubule pools (Figure 1B). It is of note that, unlike previous stages of the abscission process (Figure S1B), the MBR did not show large microtubule bundles flanking the FB (Figure 1B).

**Figure 1.**
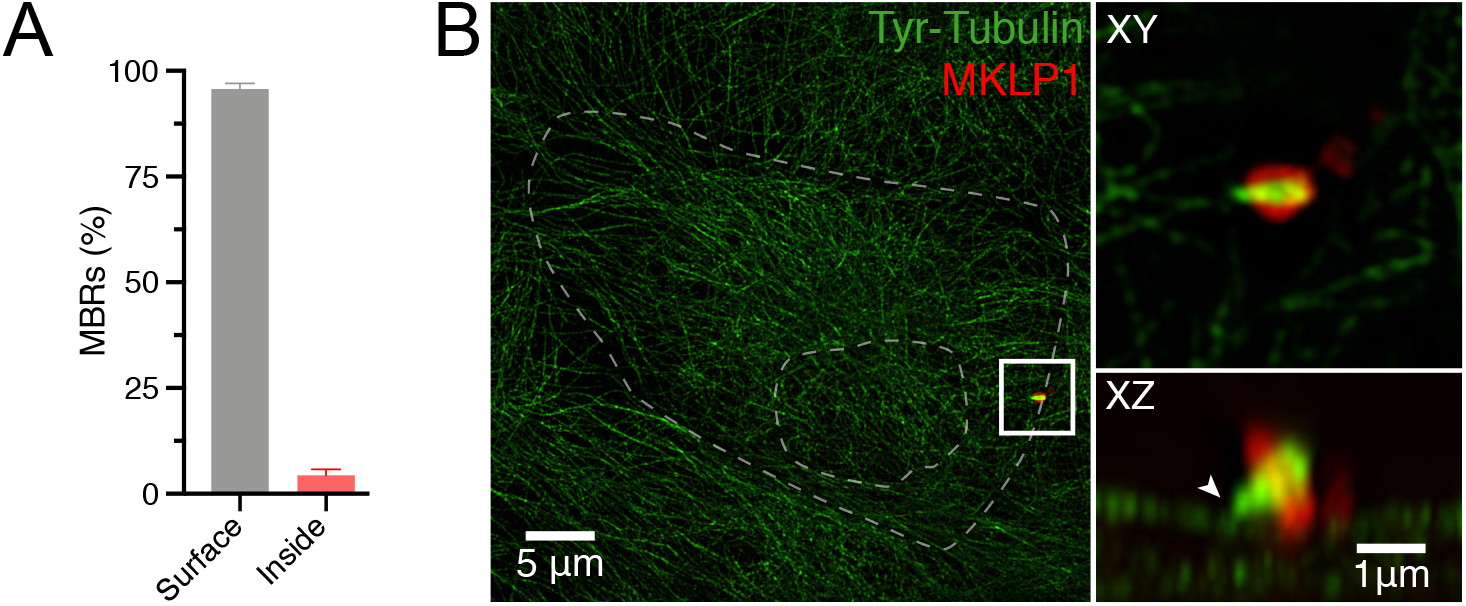
The MBR is on the surface of MDCK cells. (**A**) The localization of MBRs was analyzed by confocal microscopy. Data are summarized as the mean ± SD of the percentage of surface and intracellular MBRs from three independent experiments (n=285-296 cells). (**B**) An MBR as seen by super-resolution confocal microscopy in cells stained for tyrosinated α-tubulin and the FB marker MKLP1. Dashed lines indicate cell and nuclear contours. The enlargement of the boxed region shows the characteristic ring-like structure of the FB flanked by microtubules, as seen in both XY and XZ views. The arrowhead indicates the absence of microtubules in the region adjacent to cell body. See also Figure S1.

Abscission requires both the MB membrane and microtubules to be severed. Loss of tubulin staining on one side of the FB, coupled with the retraction of the structure, is a reliable indicator of the first membrane cleavage event. On the other side of the FB, however, additional techniques should be used to ascertain the integrity of the remaining membranous MB arm. SEM is a powerful tool for examining cell-surface topography. The most recent generation microscopes equipped with field emission tips and very-low-voltage (VLV) operation capabilities (incidence electron beam energy E_0_ ≤1 keV) allow direct, high-resolution imaging of cells on glass substrates without the need for metal coating (Wuhrer and Moran, 2016). To investigate the existence of a membranous stalk connecting the MBR membrane and the plasma membrane, we used correlative light microscopy and VLV SEM (CLEM)) in cultures of cells stably expressing GFP-tubulin (Figure S2). Light microscopy, on the one hand, allowed selection of MBR candidate structures by the strong labeling of the FB with GFP-tubulin, discarding native MBs or MB-derived structures that still maintain microtubule bundles flanking the FB. Only the structures previously selected that had typical morphology under the SEM —a bulge flanked by two cones (Chen et al., 2013, Crowell et al., 2014)— and size (1-2 μm long) were considered MBRs. Inspection of the candidate structures by VLV SEM identified unambiguously 87 *bona fide* MBRs of 117 structures analyzed. As revealed by CLEM, MBRs consist of a central “core” region, which corresponds to the bulge observed by transmission EM that contains the FB (Byers and Abramson, 1968), flanked by two opposed conical structures (Figures 2A and B). In top-view images, some of the MBRs examined have an evident membranous connection, emerging from one of the cones, with the plasma membrane (Figure 2A) that is absent from other MBRs (Figure 2B). After acquisition of a top-view image, the sample stage was tilted through 45° and rotated (Figure S2), making it possible to observe the MBR from different angles to examine the existence of a connection (Figures 2A and B, middle and right panels, and Video 1). MBRs lacking microtubular connections with the cell body and showing membrane continuity between the tip of one of the cones and the plasma membrane were classified as connected. We reasoned that the connection should restrict MBR movement in live cells in such a way that the MBR could move, defining a funnel-shaped volume whose narrowest end coincides with the connection point (Figure 2C). To confirm the existence of the connection, we carried out time-lapse analysis of MBR movement and observed that this was the case (Figures 2D and E, and Video 2). In summary, the two independent experimental approaches used support the existence of a physical connection between some MBRs and the plasma membrane.

**Figure 2.**
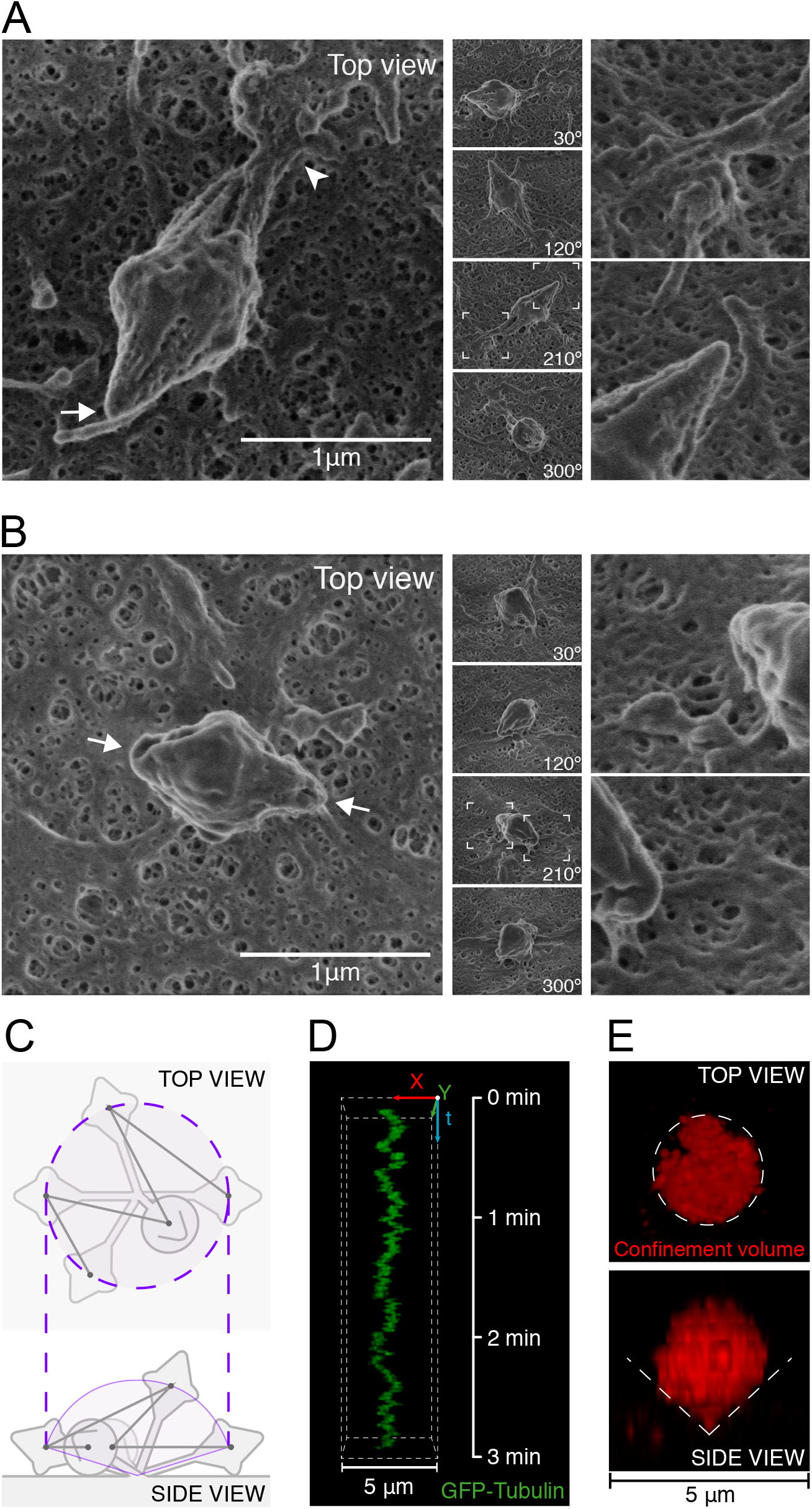
Analysis of MBRs by correlative light and scanning electron microscopy. (**A**, **B**) Images of a connected (A) and a non-connected MBR (B) on the plasma membrane as observed by SEM in top (left panels) and side views (middle panels). Numbers indicate the angle of rotation of the sample stage. The arrowhead shows the connection point with the plasma membrane and the arrows indicate the tip of the non-connected cones. The boxed regions were enlarged to show the tip of the cones in greater detail (right panels). The conical structures at the sides of the FB of these MBRs are of similar length and, therefore, they are shown as representative of symmetrical connected (A) and non-connected MBRs (B). (**C**-**E**) Graphical representation in top and side views of the predicted confinement volume in which MBR movement is restricted (C). (D) Kymograph showing a 3D reconstruction of the movement of an MBR, as visualized with GFP-tubulin, over time in a live cell. (E) Top and side views of the funnel-shaped confinement volume calculated from the same MBR. See also Figure S2 and Videos 1 and 2.

### The membranous connection extends from the tip of the largest MBR cone

The MBRs identified in our analysis were quantified and classified according to the existence of a membranous connection with the plasma membrane, the symmetry between the two cones, and the size of the cone from which the connection arises (Figure 3A). Top-view SEM images showed a clear connection with the plasma membrane in 45/87 of the MBRs, whereas no discernible connection was found in 17/87 MBRs (Figure 3B). The remaining 25/87 MBRs were classified as “unclear” because of their arrangement on the cell surface precludes the visualization of the possible connection in top-view images (Figure 3B and S3A). The number of “informative” (45 + 17) top-view images of MBRs was considered sufficient to make further analysis of unclear cases unnecessary. The inclination angle formed by the long axis of the MBR and the cell surface observed for the unclear cases was more similar to that of the clearly connected MBRs than to those of the non-connected ones (Figures S3B and C), suggesting the presence of a connection in most of the unclear cases. This observation implies that the observed fraction of connected MBRs with respect to the total “informative” cases (45/62, or 72.6%) is likely an underestimate of the genuine fraction of connected MBRs.

**Figure 3.**
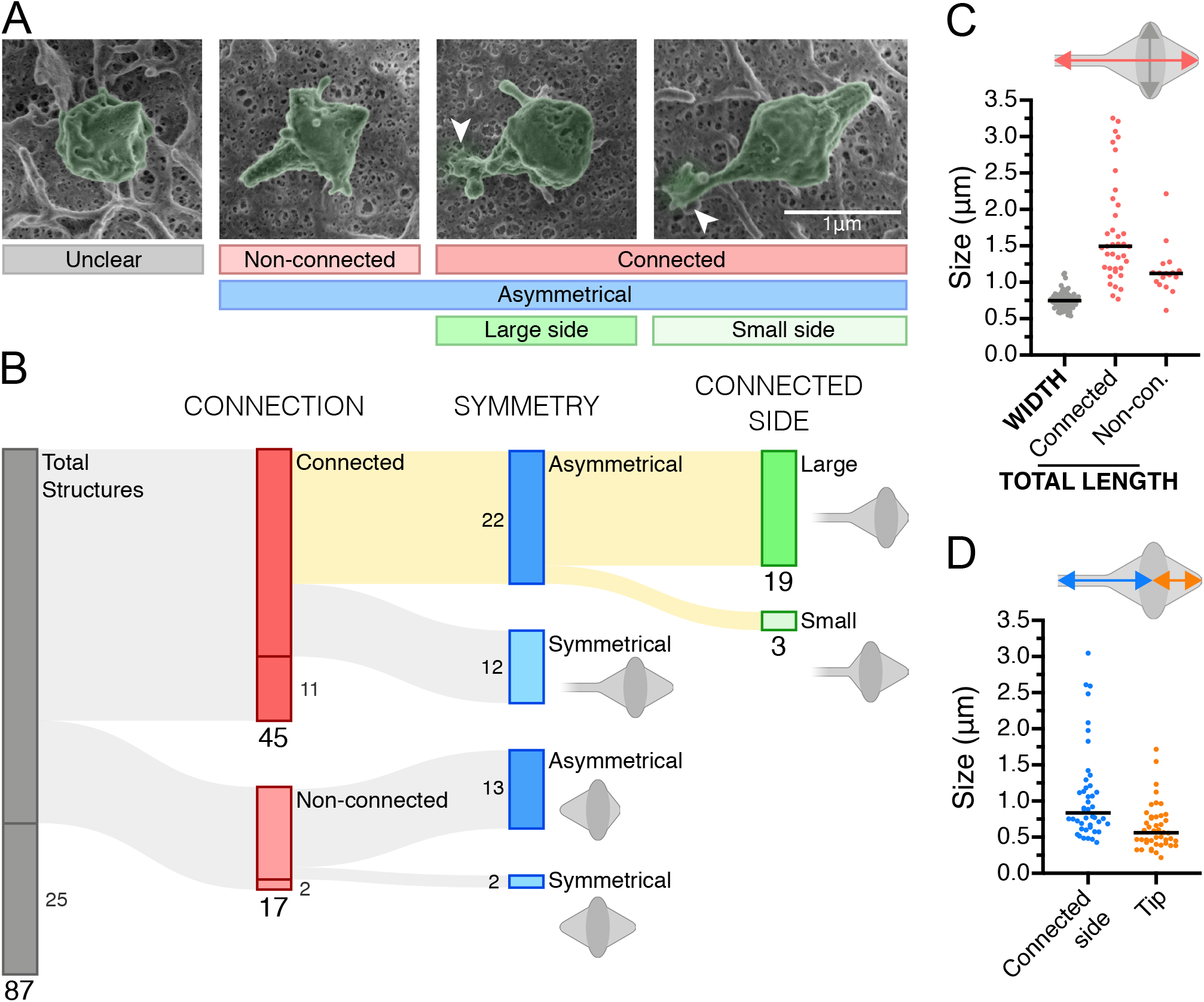
Most MBRs remain connected to the plasma membrane. (**A**) Representative examples of MBR morphologies others than those shown in Figure 2. Arrowheads indicate connection points. (**B**) Sankey diagram showing the results of our MBR morphology analysis. Large and small sized numbers indicate the population size of each class and of the subclasses, respectively. Only the structures with strong labeling of the Flemming body with GFP-tubulin, that lacked microtubular connections with the cell body on both flanks, and that had typical morphology and size were considered MBRs. (**C**) Quantification of FB width (n=87) and total MBR length of connected (n=38) and non-connected (n=17) structures. (**D**) Length of the two sides flanking the FB in connected MBRs (n=38). Black bars indicate median values. See also Figure S3.

A morphological feature of MBRs is the apparent degeneration of one of the cones. While one cone tends to have a defined form and size, the other is frequently shorter and rounder, giving rise to an asymmetrical MBR (Figure 3A). It is of note that the connection arose from the larger cone in most (19/22, or 86.4%) of the connected MBRs with asymmetrical cones (Figure 3B).

To characterize the MBR, we measured its dimensions using top-view SEM images. Connected MBRs were longer and more variable in length than the non-connected ones, being the connected side longer than the opposite one (Figures 3C and D). Independent measurements of the length based on MBR movement yielded similar values, supporting the validity of this approach (Figures S3D-F).

In conclusion, the analyses presented so far indicate that MBRs display a number of prevalent structural features, the most common one being the presence of a membranous stalk presumably derived from the uncleaved arm of the bridge, which most often coincides with the largest cone, physically connecting the MBR membrane to the plasma membrane.

### The ESCRT machinery concentrates at the connection between the MBR and the plasma membrane

The final steps of the abscission process are carried out by the ESCRT machinery (Carlton and Martin-Serrano, 2007, Morita et al., 2007, Schoneberg et al., 2017), which progressively accumulates into rings at both sides of the FB (Elia et al., 2011). ESCRT-III assembles spiral polymers whose diameter decreases as they grow away from the FB, constricting the MB to the limit allowed by the microtubules inside. After microtubule clearance, the ESCRT polymer remodels generating a second ESCRT pool that is positioned at the future cleavage site (Elia et al., 2012, Goliand et al., 2018).

To investigate the involvement of ESCRT-III proteins in the cleavage of the membrane of the other MB arm, we expressed GFP-fused forms of the ESCRT proteins CHMP4B (GFP-L-CHMP4B) and CHMP4C (GFP-L-CHMP4C), and analyzed their localization before and after the end of the abscission process. These proteins, in which GFP is separated from CHMP4C and CHMP4B by a 25-nm long flexible linker, were previously shown to have the expected localization at the MB, and their expression did not delay MB abscission time (Ventimiglia et al., 2018, Sadler et al., 2018). Both proteins first accumulated in ring-like structures on both sides of the FB and then polymerized towards the abscission site, resulting in the appearance of cone-shaped staining in one of the MB arms. Once microtubules were cleared from this arm, membrane cleavage and, consequently, daughter cell separation occurred. After abscission, CHMP4B, CHMP4C and microtubules followed essentially the same sequence of events on the other side of the FB, generating an MBR (Figures 4A and B, and Figure S4A). The same was observed in a panel of endogenous ESCRT proteins (Figure S4B) and by videomicroscopic analysis of cells co-expressing GFP-L-CHMP4B and Cherry-tubulin (Video 3). All MBRs contained ESCRT proteins (Figure S4C) but the pattern of distribution was not the same in all MBRs. The MBRs that presented a similar pattern on both sides of the FB, mainly with staining only on the FB rims, were classified as “even” MBRs, whereas those that, in addition to the FB rims, had a second ESCRT pool in only one side of the FB were categorized as “uneven” MBRs. The second ESCRT pool in the uneven MBRs adopted the form of a cone, filament or dot (Figure 4A and B, Figure S4D, and Video 3). Quantitative analysis revealed that most MBRs display uneven ESCRT distribution (Figure 4C). It is of particular note that the MBR side with the extra ESCRT pool coincides with that having the membranous stalk. This pool is present in a region of the connection proximal to the plasma membrane, as determined by CLEM of cells stably expressing Cherry-tubulin and GFP-L-CHMP4C or GFP-L-CHMP4B (Figure 5A-E, and Videos 4 and5). Supporting this localization, time-lapse analysis of MBR movement showed that the pool remained immobile, as may be seen in the projected kymograph, whereas the distal pool, which corresponds to the FB rims, moved drawing a circle around it (Figure 5F and Video 6).

**Figure 4.**
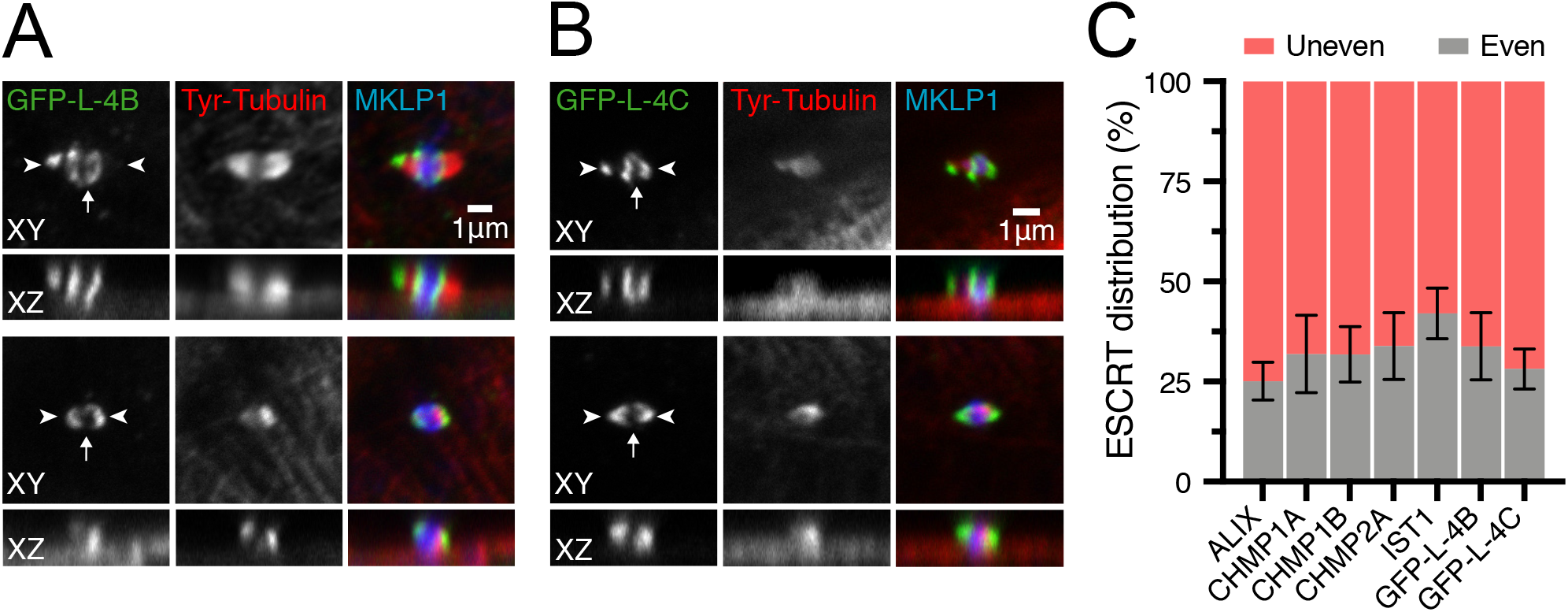
The ESCRT machinery shows an uneven distribution along most MBRs. (**A**, **B**) Distribution of GFP-L-CHMP4B (GFP-L-4B) (A) and GFP-L-CHMP4C (GFP-L-4C) (B) at the MBR. XY and XZ views of MBRs with uneven (top panels) and even (bottom panels) distribution of these markers. The arrow and the arrowheads in A and B indicate the FB and the MBR tips, respectively. (**C**) Histogram showing the percentage of MBRs with uneven and even distribution for GFP-L-CHMP4B, GFP-L-CHMP4C and a panel of endogenous ESCRT markers. Data are summarized as the mean ± SD from three independent experiments (n=29-93). See also Figure S4 and Video 3.

**Figure 5.**
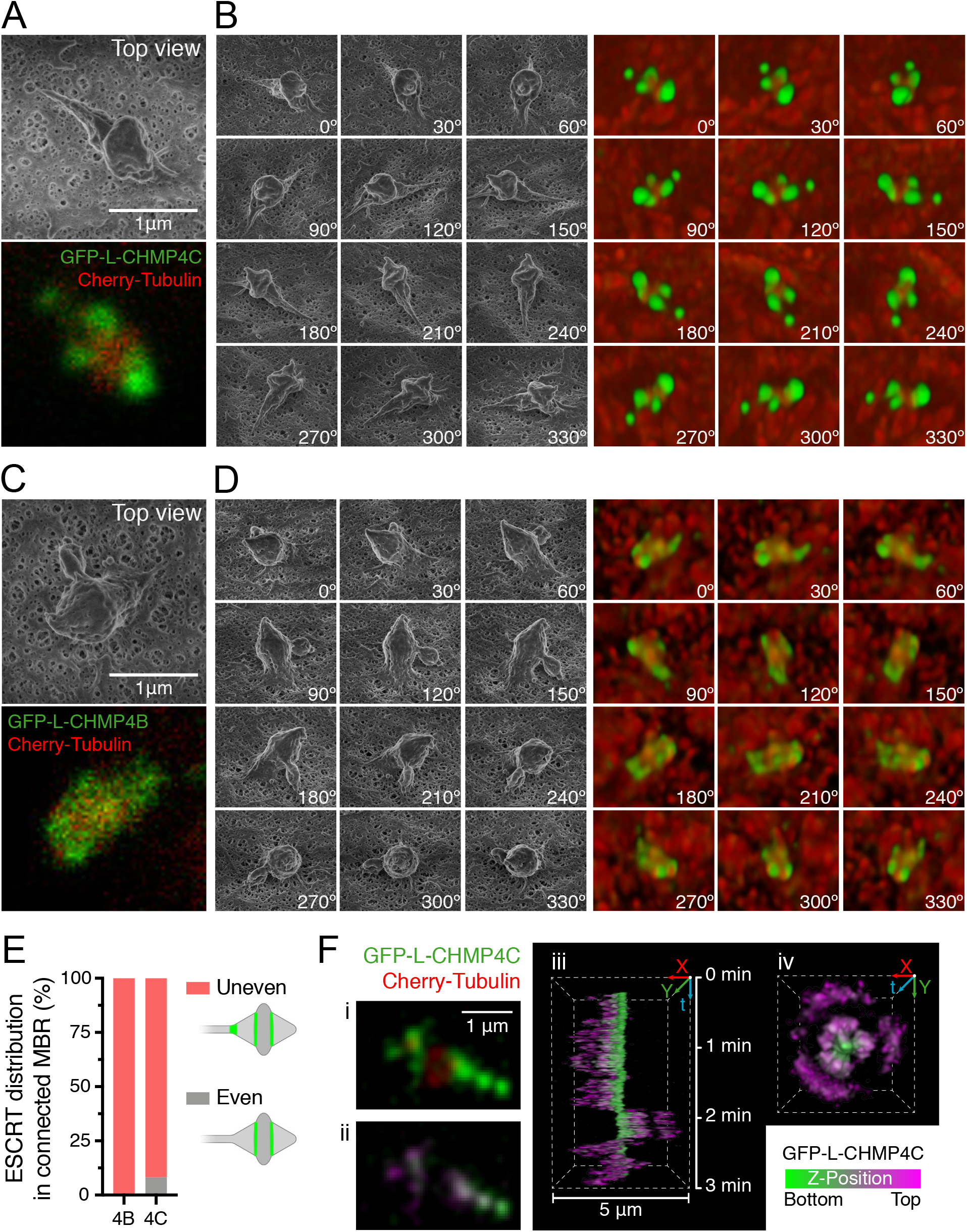
CHMP4C and CHMP4B are present at the membranous connection. (**A**-**D**) CLEM images showing the presence of GFP-L-CHMP4C (A, B) and GFP-L-CHMP4B (C, D) at the connection of the MBR with the plasma membrane. (A, C) top-view images of connected MBRs acquired by SEM (top) and confocal microscopy (bottom). (B, D) Side-view SEM images (left) and matching confocal images obtained by 3D reconstruction (right). (**E**) Quantification of GFP-L-4B and GFP-L-4C distribution in connected MBRs as observed by CLEM (n=16 and 18, respectively). (**F**) Tracks of GFP-L-CHMP4C and Cherry-tubulin movement of an MBR in a live cell. (i) GFP-L-CHMP4C and Cherry-tubulin distribution in an MBR; (ii) image of the distribution GFP-L-CHMP4C using the indicated depth-color scale; (iii and iv) 3D reconstruction of the movement followed by the MBR over a 3-min period. See also Videos 4-6.

In summary, ESCRT proteins localize to the membranous stalk that connects the MBR to the plasma membrane and have a similar distribution to that found in pre-abscission stages right before the MB arm is first cleaved (Goliand et al., 2018). Since the presence of an ESCRT pool distant from that surrounding the FB has been associated with the last stage of membrane cleavage (Goliand et al., 2018), we proceeded to analyze how the cleavage of the connection is prevented.

### CHMP4C depletion reduces the percentage of cells with an MBR and impairs primary ciliogenesis

The abscission checkpoint delays abscission by regulating the ESCRT machinery in the case of mitotic problems, such as persisting chromatin within the bridge, incomplete nuclear pore reformation defects, or tension in the bridge produced by opposite pulling forces from the daughter cells (Agromayor and Martin-Serrano, 2013, Caballe et al., 2015). The activation of the abscission checkpoint retards abscission by promoting the phosphorylation of the ESCRT-III subunit CHMP4C by the kinase Aurora B, Ser210 being the major phospho-acceptor residue (Carlton et al., 2012, Capalbo et al., 2012). To investigate the involvement of this mechanism in the regulation of the second cleavage of the MB, we used specific siRNA (siCHMP4C) to knockdown (KD) CHMP4C expression (Figure 6A and B). This siRNA targets the sequence in dog CHMP4C equivalent to that of the siRNA previously used for human CHMP4C in HeLa cells (Carlton et al., 2012). As a control, we observed that CHMP4C KD accelerated abscission (Figure 6C) without affecting the number of dividing cells (Figure S5A), as has been noted in other cell lines (Carlton et al., 2012, Sadler et al., 2018, Caballe et al., 2015). The acceleration of abscission is statistically significant but modest, since it only affects the minority of cell divisions in which abscission is delayed by the abscission checkpoint mechanism. It is of note that the percentage of cells with an MBR was much lower in CHMP4C KD cells that in the control cells (Figure 6D). Videomicroscopic analysis showed MBR shedding in CHMP4C KD cells (Video 7). As a control, we observed that CHMP2A KD did not affect the percentage of cells with an MBR (Figure 6D and Figures S5B and C), suggesting that there is a specific effect of the regulatory CHMP4C subunit, but not of any ESCRT component, in the regulation of MBR inheritance. The CHMP4C mutants S210A and A232T, which is a CHMP4C allele associated with increased susceptibility to cancer, are unable to replace endogenous CHMP4C in the regulation of the first cut of the bridge membrane (Carlton et al., 2012, Sadler et al., 2018). The effect of CHMP4C knockdown on the second cut was rescued by the exogenous expression of siCHMP4C-resistant forms of GFP fusions of wild type but not of the S210A and A232T CHMP4C mutants (Figure 6D, and Figures S5D-F). The percentage of MBRs positive for the CHMP4C mutants, the distribution of the mutants within the MBR, and the total number of cells per field were similar to those of the wild-type CHMP4C protein (Figures S5G-I). As a control, we observed that the number of cells connected by a MB decreased in siCHMP4C-treated cells and that this effect was corrected by the intact protein but not by the CHMP4C mutants (Figure S5J). The results illustrated in Figure 6C for the second cut of the membrane of the bridge are consistent with those reported for CHMP4C in the control of the first cut by the abscission checkpoint mechanism (Carlton et al., 2012, Capalbo et al., 2012). They suggest that CHMP4C has a similar role in the second cut.

**Figure 6.**
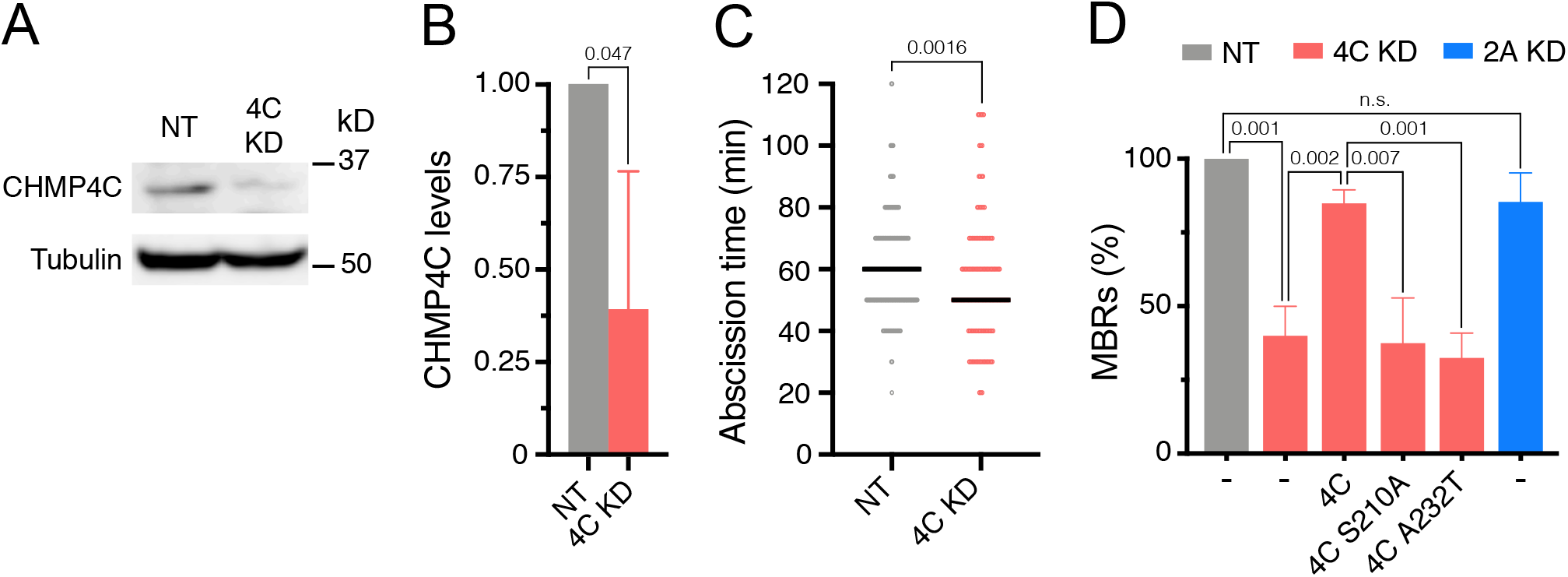
CHMP4C is required for MBR inheritance. (**A**) Representative immunoblot showing the effect of siCHMP4C on endogenous CHMP4C levels. (**B**) Quantification of CHMP4C KD. The histogram represents the levels of CHMP4C in siCHMP4C-transfected cells relative to that of cells transfected with control siNT. (**C**) The time between the formation of the MB and abscission was measured in control siNT (gray points) and siRNA-mediated CHMP4C KD cells (red points) (n=27-159 in control cells; n=8-95 in CHMP4C KD cells). Black bars represent median values. (**D**) The percentage of MBRs ± SD in CHMP4C KD cells (red bars) expressing the indicated exogenous CHMP4C proteins and of CHMP2A KD cells (blue bar) was calculated relative to that of control cells (siNT, gray bars) (n=2714-3447 cells for control and n=800-1463 for KD cells). Three independent experiments were performed. Probabilities are those associated unpaired two-tailed Student’s t-test (B, D) and Mann-Whitney test (C). See also Figure S5 and Video 7.

Most mammalian cells have a primary cilium that projects from their surface as a single appendage (Wheatley et al., 1996). The primary cilium orchestrates important signaling pathways involved in development and cell proliferation, differentiation, survival and migration (Malicki and Johnson, 2017, Goetz and Anderson, 2010). Primary cilium formation proceeds by two distinct pathways, depending whether the position of the centrosome in the cell (Sorokin, 1968). In cells with centrosome near the nucleus, as in NIH 3T3 fibroblasts, ciliogenesis starts intracellularly and finishes at the plasma membrane, whereas when the centrosome is close to the plasma membrane, as in polarized epithelial MDCK cells, the process takes place entirely at the plasma membrane. The first route is referred to as the intracellular or “classic” pathway; the second route is known as the alternative pathway (Albrecht-Buehler and Bushnell, 1980, Molla-Herman et al., 2010, Bernabe-Rubio and Alonso, 2017). Since we described previously that the MBR plays an important role in preparing the centrosome for primary ciliogenesisin MDCK cells (Bernabe-Rubio et al., 2016), we examined the effect of CHMP4C KD on this process. We noted a dramatic drop in the percentage of ciliated cells (Figures 7A and B), which is consistent with the loss of MBRs in CHMP4C-deficient cells (Figure 6D). Confirming the requirement for CHMP4C, the exogenous expression of siCHMP4C-resistant wild type CHMP4C rescued primary cilium formation (Figures 7A and B). The results in Figure 7 are in agreement with a previous report showing that the physical removal of the MBR greatly reduces primary ciliogenesis (Bernabe-Rubio et al., 2016) and further highlights the importance of the MBR in this process by providing a genetic evidence of the requirement for MBR in primary cilium formation by polarized epithelial cells.

**Figure 7.**
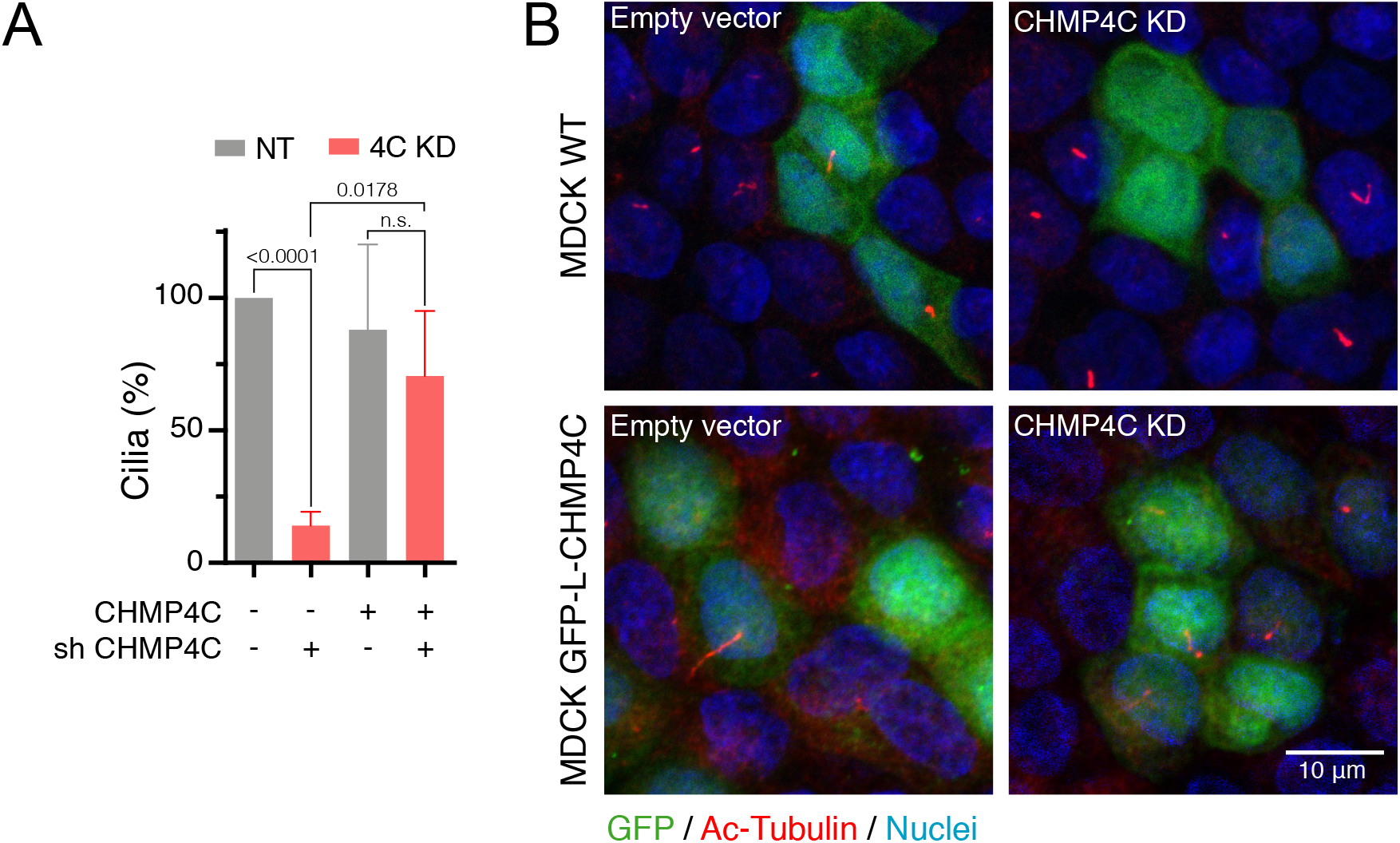
CHMP4C is required for primary cilium formation. (**A**) Effect of CHMP4C KD on the frequency of ciliated cells. The percentage of primary cilia in cells expressing shCHMP4C and GFP in the absence or presence of exogenous human CHMP4C was expressed relative to that of control cells, which expressed only GFP (n=77-88 for control; n=70-161 for CHMP4C KD cells). The mean ± SD from three independent experiments is show. Probabilities are those associated with unpaired two-tailed Student’s t-test. (**B**) Representative fields of the cells analyzed in (A). Acetylated tubulin staining was used to visualize the primary cilium.

## Discussion

Although the FB was first described more than 125 years ago, the discovery of its role in abscission is relatively recent, and even more so is the evidence of important post-mitotic roles for the MBR (Chen et al., 2013). Accumulation of MBRs has been associated with increased cell reprogramming efficiency of stem cells and *in vitro* tumorigenicity of cancer cells (Kuo et al., 2011, Ettinger et al., 2011, Peterman et al., 2019). In polarized epithelial cells, the MBR meets the centrosome at the center of the apical membrane and enables the centrosome for primary cilium formation (Bernabe-Rubio et al., 2016). Using CLEM, we identified a membranous stalk in polarized epithelial MDCK cells that physically connects the MBR membrane and the plasma membranes of most MBR-containing cells. The stalk is derived from the uncleaved arm of the bridge and contains ESCRT machinery, including the regulatory subunit CHMP4C. CHMP4C silencing causes the loss of the MBR and, consistent with its role in primary cilium formation, a dramatic reduction in the percentage of ciliated cells. These results indicate that an MBR physically connected to the plasma membrane by a membranous stalk, whose integrity is regulated by CHMP4C, is the form of MBR used by MDCK cells to license primary ciliogenesis.

We first identified candidate MBR structures from the presence of GFP-tubulin in the MB core and its absence from the two MB arms. The selected structures were analyzed in a state-of-the-art, VLV SEM using samples that were prepared by a gentle procedure (Katsen-Globa et al., 2016) omitting conductive coating. This equipment revealed the subnanometric topography of MBRs, which enabled structures without the typical MBR morphology to be discounted. Using this approach, we visualized a membranous stalk between the MBR and the plasma membrane in a large proportion of MBRs. However, such a connection was not observed in a previous CLEM study (Crowell et al., 2014) that combined phase-contrast microscopy to identify MBR candidates, sample preparation by standard procedures, and analysis under conventional SEM equipment (Fremont and Echard, 2017). The discrepancy between the two studies might be due to the different cell lines analyzed —HeLa cells in Crowell et al. (2014) and MDCK cells in ours— or to the distinct protocols for sample preparation and the SEM equipments used. In addition to detecting the connection, our CLEM analysis revealed that one of the MBR cones is larger than the other, likely because the shorter one results from the degeneration of the cone on the side where abscission occurs. Consistent with this possibility, we observed that the connecting stalk most often arises from the largest cone of the MBR. The degeneration of the connected cone is probably prevented by the presence of both conical ESCRT polymers (Sherman et al., 2016), which could act as a scaffold for the structure, and the microtubules filling the cone.

The loss of tubulin staining on one side of the FB together with the retraction of the structure indicates the first cut of the membrane of intercellular bridge (Elia et al., 2011, Lafaurie-Janvore et al., 2013). However, it is important to note that the loss of tubulin staining on the other side should not be assumed to represent the second cut of the bridge membrane, since CLEM revealed the existence of a membranous connection with the plasma membrane in MBRs with no flanking microtubules. In addition, the use of phase-contrast microscopy is not adequate for distinguishing MBRs that are attached to the plasma membrane from those that are physically connected by a membranous stalk because the connection is very small (Crowell et al., 2014). Therefore, our work is consistent with previous studies (Elia et al., 2011, Lafaurie-Janvore et al., 2013), but differs in that we have investigated whether the second cut of the bridge membrane takes place, and have used CLEM as a highly reliable technical approach to do precisely so.

We observed that most MBRs contained ESCRT polymers only on the side corresponding to the largest cone, similar to those present just before the first cleavage of the MB membrane. We mapped the ESCRT pool at the membranous connection between the MBR and the plasma membrane by CLEM, and confirmed the localization by analyzing the MBR motion. This location of ESCRT proteins is consistent with the presence of helical filaments in the uncleaved MB arm, as observed by soft X-ray cryotomography (Sherman et al., 2016). This pool contains CHMP4C, which is a crucial component of the checkpoint mechanism that delays abscission when mitotic problems occur. In those cases, the knockdown of CHMP4C accelerates abscission and only the expression of wild type CHMP4C but not of the CHMP4C S210A or A232T mutants can substitute the endogenous protein to delay membrane cleavage. Since the number of cells with an MBR was greatly diminished in CHMP4C KD cells and the effect was corrected by expression of intact CHMP4C but not by CHMP4C mutants, we propose that, similar to its role in the abscission checkpoint (Carlton et al., 2012, Capalbo et al., 2012), CHMP4C allows MBRs to remain connected to the plasma membrane by delaying the cleavage of the connection.

The model of primary ciliogenesis based on MBR-mediated centrosome licensing that we proposed implies that only cells with an MBR can assemble a cilium (Bernabe-Rubio et al., 2016). Therefore, MBR retention is important at the cell-population level to guarantee a high percentage of MBR-bearing cells, and at the single-cell level to permit the centrosome to form a primary cilium. The existence of the physical connection in MDCK cells might facilitate, at the cell population-level, the retention of the MBR to be transmitted to the next generation of cells, as seen by videomicroscopic analysis (Bernabe-Rubio et al., 2016), and, at the single-cell level, the directional movement of the MBR to meet the centrosome by direct anchoring to the cytoskeleton. In addition, the continuity of the MBR with the rest of the plasma membrane might enable the transfer of materials from the MBR to the centrosome in order to assemble the primary cilium. There are conflicting results about the requirement of ESCRT proteins for primary ciliogenesis in cells such as NIH 3T3 cells that, unlike MDCK cells, use the intracellular route of ciliogenesis. Primary cilium formation appears to be independent of ESCRT proteins in one of the studies (Ott et al., 2018) but not in the other (Jung et al., 2020). Since we found that a functional consequence of the loss of the connection caused by CHMP4C silencing is the impairment of primary ciliogenesis in MDCK cells, we conclude that the connection is required for the alternative route of primary ciliogenesis.

In conclusion, our study reveals that the majority of MBRs inherited in MDCK cells are physically connected to the plasma membrane through a membranous stalk derived from the uncleaved arm of the cytokinetic bridge. The ESCRT subunit CHMP4C controls the integrity of this arm to ensure the continuity between the MBR membrane and the plasma membrane and, in this way, the MBR facilitates primary cilium formation.

## Materials and Methods

### Antibodies

The sources of the antibodies to the different markers were as follows: total α-tubulin (mouse mAb IgG1, clone DM1A, product T6199; used at 1/5,000), tyrosinated α-tubulin (rat mAb IgG2a, clone YL1/2, product MAB1864; used at 1/200), acetylated tubulin (mouse mAb IgG2b; clone 6-11-B1, product T7451; used at 1/500) and CHMP1B (rabbit polyclonal antibody, ATLAS product HPA061997; used at 1/500) were from Merck; CHMP2A (rabbit polyclonal, product 10477-1-AP; used at 1/500 for immunofluorescence analysis and at 1/000 for western blot) was from Proteintech; CHMP1A (rabbit polyclonal, product ab178686; used at 1/500) was from Abcam; PRC1 (mouse mAb IgG2b, clone 16F2, product MA1-846; used at 1/100) was from ThermoFisher Scientific; MKLP1 (rabbit polyclonal, product sc-867; used at 1/100) was from Santa Cruz; GFP (mouse mAbs IgGκ, mixture of clones 7.1 and 13.1, product 11814460001; used at 1:1,000) was from Roche. The rabbit polyclonal antibody to CHMP4C was prepared by Lampire Biologicals and used at 1/200. The rabbit polyclonal antibodies to ALIX (used at 1/500) and IST1 (used at 1/1,000) (Bajorek et al., 2009) were generous gifts from Dr. Sundquist (University of Utah). Secondary antibodies conjugated to Alexa-488, −555 or −647 were from Thermo Fisher Scientific.

### Cell culture

Epithelial canine MDCK II (CRL2936) cells were obtained from the ATCC and grown in MEM supplemented with 5% FBS (Merck) at 37°C in an atmosphere of 5% CO_2_. Mycoplasma testing was regularly performed. For immunofluorescence and quantitative analysis, 3.0×10^4^ cells were plated onto coverslips maintained in 24-well multiwell plates and grown for 48 h. For correlative light and electron microscopy and time-lapse studies 1.5×10^5^ cells were plated onto 35-mm glass-bottom plates (MatTek) and grown for 48 h.

### DNA constructs, siRNA and transfection conditions

The DNA constructs expressing EGFP- or mCherry-tubulin were from Takara Bio, Inc. MDCK II cells stably expressing these proteins were generated by transfection of 1.0×10^6^ cells with Amaxa Nucleofector II (Lonza) using the L-005 program. After selection with 2 mg/ml G-418 (Thermo Fisher Scientific), the resulting clones were screened under a fluorescence microscope. The retroviral constructs pNG72-GFP-L-CHMP4B, pNG72-GFP-L-CHMP4C, pNG72-GFP-L-CHMP4C A232T have been described previously (Ventimiglia et al., 2018, Sadler et al., 2018). pNG72-GFP-L-CHMP4C S210A was generated by site-directed mutagenesis using a commercial kit (Quickchange Lightning, Agilent Technologies). For retroviral production, 293T cells were co-transfected with the indicated retroviral construct and with the retroviral packaging vectors, MLV-GagPol/pHIV 8.1 and pHIT VSVg at a ratio of 2:3:1 for 48 h using polyethylenimine (Polysciences, Germany). 293T supernatant was collected and filtered through a 0.2-μm filter before being used to transduce MDCK II cells. For siRNA assays, 3.0×10^4^ cells were transfected with 100 nM (ThermoFisher Scientific) non-targeting siRNA (siNT), custom siRNA targeted to dog CHMP4C (siCHMP4C, 5’-CTCGCTCAGATTGATGGCACA-3’) (Carlton et al., 2012) or a 1:1 mixture of two siRNA to CHMP2A (5’-CAGGCCGAGAUCAUGGACAUG-3’ and 5’-GAAGAUGAAGAGGAGAGUGAC-3’) using Lipofectamine 2000 (Thermo Fisher Scientific) according to the manufacturer’s recommendations. Cells were transfected twice, 48 h and 6 h before the beginning of the experiments. The pSuperGFP-shCHMP4C construct, which expresses GFP and shRNA to dog CHMP4C simultaneously from independent promoters, was generated by cloning a synthetic DNA duplex with the same target sequence as siCHMP4C into the pSuper plasmid (OligoEngine). The resulting plasmid was combined with the plasmid pEGFP-N1 using the unique EcoO109I and Afl III sites present in both plasmids.

### Confocal microscopy

Cells were fixed in cold methanol for 5 min. After blocking with 3% (wt/vol) BSA for 30 min, cells were incubated with the indicated primary antibodies at 4°C overnight, and were washed and then stained with the appropriate fluorescent secondary antibodies. Coverslips were mounted using ProLong Gold antifade reagent (ThermoFisher Scientific). Super-resolution images were obtained using a Nikon N-SIM-S superresolution microscope with a 100x oil immersion objective (Numerical aperture, NA, of 1.49) and processed with NIS-Elements. A stack containing the whole cell was acquired in 3D-SIM imaging mode. Maximum intensity projections of the entire stack are shown. Images for ESCRT localization analysis were acquired with an LSM 800 confocal microscope (Zeiss) equipped with a 63x oil immersion objective (NA 1.4) and a Nikon Eclipse Ti-E inverted CSU-X1 spinning disk confocal microscope equipped with a 100x oil immersion objective (NA 1.4). The images shown are the sums of the planes containing the structure of interest. The surface or intracellular localization of MBRs was analyzed by confocal microscopy using XZ projections of 18 planes.

### Time-lapse confocal imaging

Cells were seeded on 35-mm glass-bottom dishes as mentioned above and maintained in MEM without phenol red during recording. Time-lapse experiments showing MBR motion were acquired with a Nikon A1R+ confocal microscope with a 60x water objective (NA 1.2). A stack containing the whole structure was captured every second using a resonant scanner, and the resulting images were deconvoluted with Huygens software (SVI) to enhance the signal-to-noise ratio. 3D reconstructions were generated in NIS-Elements software (Nikon). To quantify motion confinement, a single plane was acquired every second for 3 min with a Nikon Eclipse Ti-E inverted CSU-X1 spinning disk confocal microscope equipped with a 100x oil immersion objective (NA 1.4). The position of the structure was determined in every frame, and the geometrical center of every dataset calculated. From that point, a circle that included 95% of the points was delineated and used to calculate the length of the MBR connection.

### Correlative light and scanning electron microscopy

Prior to cell seeding, 250-nm gold nanobeads (BBI Solutions) were deposited over a 35-mm glass-bottom plate pre-coated with polylysine (1.0×10^4^ beads/mm^2^) to serve as fiducial markers. Reference marks were made on the coverslip to localize the imaging area and maintain sample orientation between the two imaging methods used. Then, MDCK cells stably expressing GFP-tubulin alone or Cherry-tubulin plus either GFP-L-CHMP4B or GFP-L-CHMP4C were seeded as described above. After 48 h of cell growth, cells were pre-fixed with a volume of 2x fixing solution (4% paraformaldehyde plus 4% glutaraldehyde in phosphate buffer) equal to that of the culture medium for 10 min at room temperature, followed by 3 h incubation with 1x fixing solution. For the confocal microscopy component, a Nikon A1R+ confocal microscope with a 60x water objective (NA 1.2) was used. First, a low-magnification image was acquired for alignment and navigation purposes, including the fluorescence signal and a reflection channel showing the position of the gold nanobeads. Candidate MBR structures selected by the absence of tubulin label at the FB sides were identified and high-resolution images were acquired when needed. The samples for SEM analysis were prepared by a gentle procedure adapted from Katsen-Globa et al. (2016) that avoids conventional treatments, such as osmium post-fixation, critical-point desiccation, and sputter coating with gold, that could alter the cell-surface topography and that are used in sample preparation for analysis under conventional SEM equipments. Briefly, the cells in the coverslisp were dehydrated by immersion in increasing concentrations of ethanol (10% increments up to 100%, 3 min per solution). After dehydration, ethanol was substituted by hexamethyldisilazane (HMDS, Sigma-Aldrich) by sequential 3 min incubation in a 1:1 ethanol-HMDS solution and pure HMDS. The samples were air-dried overnight. Then, the coverslip was attached to a sample holder with carbon adhesive tape and encircled with copper foil to reduce charge accumulation. Scanning electron microscopy images were acquired with ultra-high resolution FEI Verios 460 field-emission SEM equipment with a calibrated resolution below 0.6 nm at 1 keV landing energy. This equipment allows obtaining more surface detail, creating less beam damage, and reducing charging effects compared with conventional SEM equipments. Sample orientation was first adjusted using the in-chamber camera and reference marks, and the imaging area localized. A low-magnification image matching the one acquired under the confocal microscope was acquired, and the position of the gold nanobeads identified. The pattern formed by the cells over the substrate was first used for rough alignment, and the position of the gold nanobeads was then used to refine the alignment, facilitating the identification of the structures of interest. VLV SEM images of the selected structures were acquired at 1 keV with a current of 13 pA by an in-lens secondary electron detector. Only the structures with strong labeling of the Flemming body with GFP-tubulin, that lacked microtubular connections with the cell body on both flanks, and that had the typical morphology and size were considered MBRs. Those with membrane continuity between the tip of one of the cones and the plasma membrane were considered connected. In these cases, the top-view SEM image shows a membranous tether arising from the MBR tip that fuses with the surface of the plasma membrane (Figure 2A). In non-connected MBRs, the lack of continuity results in the appearance of a white border around the structure that is caused by the increased emission of secondary electrons arising from the interaction of the electron beam with a sharp edge, which thereby denotes the lack of continuity (Figure 2B). To observe the structure of interest from different angles, the sample stage was tilted through 45° and rotated in 30° increments (Figure S2C). Finally, 3D reconstructions of the corresponding confocal images were generated in NIS-Elements (Nikon) and rotated to match the orientation of their corresponding SEM counterparts.

### MBR characterization and size analysis

A top-view SEM image was acquired for every candidate structure identified as an MBR by CLEM. MBRs showing continuity between the plasma membrane and the end of one of the cones flanking the MB were classified as connected MBRs. For symmetry analysis, the overall size of both regions flanking the FB was considered. The actual length of the MBR long axis was calculated from distance and angle measurements taken from top-view images (Figure 3—figure supplemental 1B) as follows:

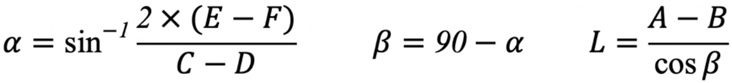

### Abscission timing quantification

Cells were seeded on glass-bottom 24-well plates (MatTek) and transfected with siRNA as previously described. Imaging was carried out with a 40x dry objective lens (NA 0.75) on a Nikon Ti-Eclipse wide-field inverted microscope controlled by NIS-Elements software (Nikon). Cells were kept at 37°C and 5% CO_2_ in an environmental chamber and imaged every 10 min for 24 h. The time period between the formation of the MB and abscission was considered as the abscission time.

### Immunoprecipitation and immunoblotting

MDCK II cells stably expressing GFP-L-CHMP4C variants were lysed at 4°C in 1 ml lysis buffer containing 50 mM Tris, pH 7.4, 150 mM NaCl, 5 mM EDTA, 5% glycerol, 1% Triton X-100 and a protease inhibitor cocktail (Merck, product 11697498001). Lysates were sonicated and centrifuged for 10 min, the cleared supernatant was then incubated with anti-GFP coupled magnetic microparticles (GFP-Trap, ChromoTek) for 2 h followed by four washing steps. Bound proteins were eluted in Laemmli’s buffer and boiled before SDS-PAGE and immunoblotting.

### Ciliogenesis assay

Cells were transfected with the plasmid (pSuperGFP-shCHMP4C) or an empty vector (pSuperGFPN1) using Amaxa nucleofector. 9.0×10^5^ cells were plated on 12-mm Transwell permeable supports (Corning) and cultured for 72 h. Samples were processed for immunofluorescence analysis and imaged with a Zeiss LSM510 confocal microscope equipped with a 63x oil immersion lens (NA 1.4). The percentages of ciliated cells were determined for the cells positive for GFP expression.

### Statistical analysis

All graphs were produced and statistical analysis performed with Prism software (GraphPad). Statistical significance was assessed with two-tailed Student’s unpaired t-test or Mann-Whitney test, as indicated. Additional information is shown in figure legends.

## Abbreviations

CHMP: charged multivesicular body protein
CLEM: correlative light and electron microscopy
ESCRT: endosomal sorting complex required for transport
FB: Flemming body
MB: midbody
MBR: MB remnant
SEM: scanning electron microscopy
VLV: very-low-voltage

## ACKNOWLEDGEMENTS

The expert technical advice of the Optical and Confocal Microscopy Facility of CBMSO is gratefully acknowledged. We thank Dr Phil Mason for revising the English language of the manuscript, Laura Fernandez-Martín for invaluable technical help and David Esteban-Mendoza for his contribution to optimization of the SEM protocol. This work was supported by a grant (PGC2018-095643-B-I00) to MAA from the Spanish Ministerio de Ciencia e Innovación (MICIN), Agencia Estatal de Investigación, y Fondo Europeo de Desarrollo Regional, European Union (MICIN/AEI/FEDER, UE), and by Wellcome Trust funding (WT102871MA) to JM-S. We also acknowledge the Micro and Nanofabrication Laboratory of the Instituto de Micro y Nanotecnología (MiNa), which is funded by the Comunidad de Madrid (S2018/NMT-4291 TEC2SPACE), MICIN (project CSIC13-4E-1794) and EU (FEDER, FSE), for invaluable help on SEM. A contract (FPU14/00295) and a short-term fellowship from EMBO to JC-A are also acknowledged. The authors declare no competing interests.

---

**Figure S1.**
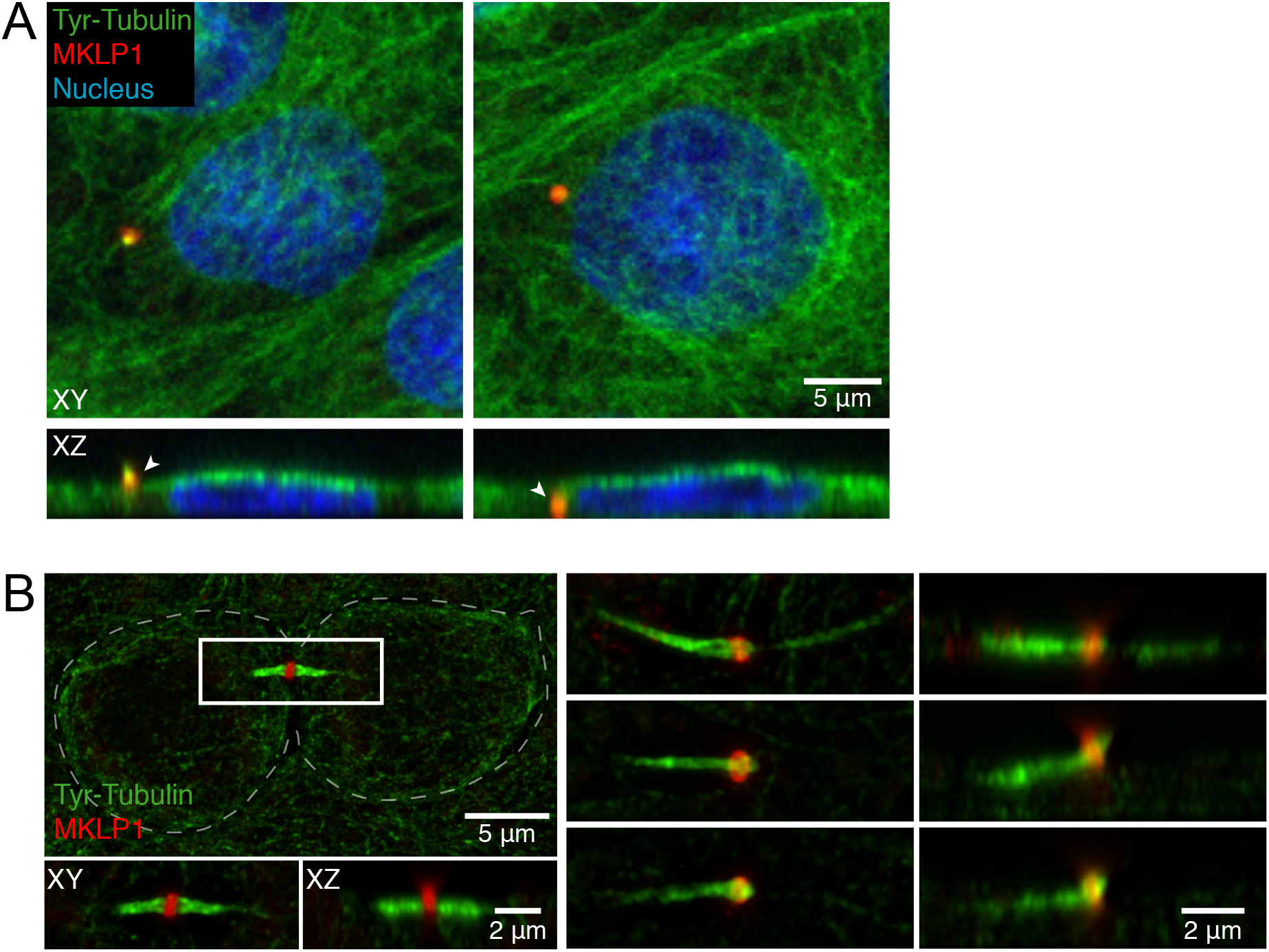
MBR localization and abscission stages in MDCK cells. (**A**) Representative examples of an MBR on the cell surface (left panels) or inside the cell (right panels). XY (top panels) and XZ projections (bottom panels) are shown. The arrowheads indicate the MBR. (**B**) Super-resolution panoramic view of two sister cells connected by an MB (left panel) and enlargement of the boxed region containing the intercellular bridge before and immediately after abscission (right panels). The dashed line delineates the cell contour.

**Figure S2.**
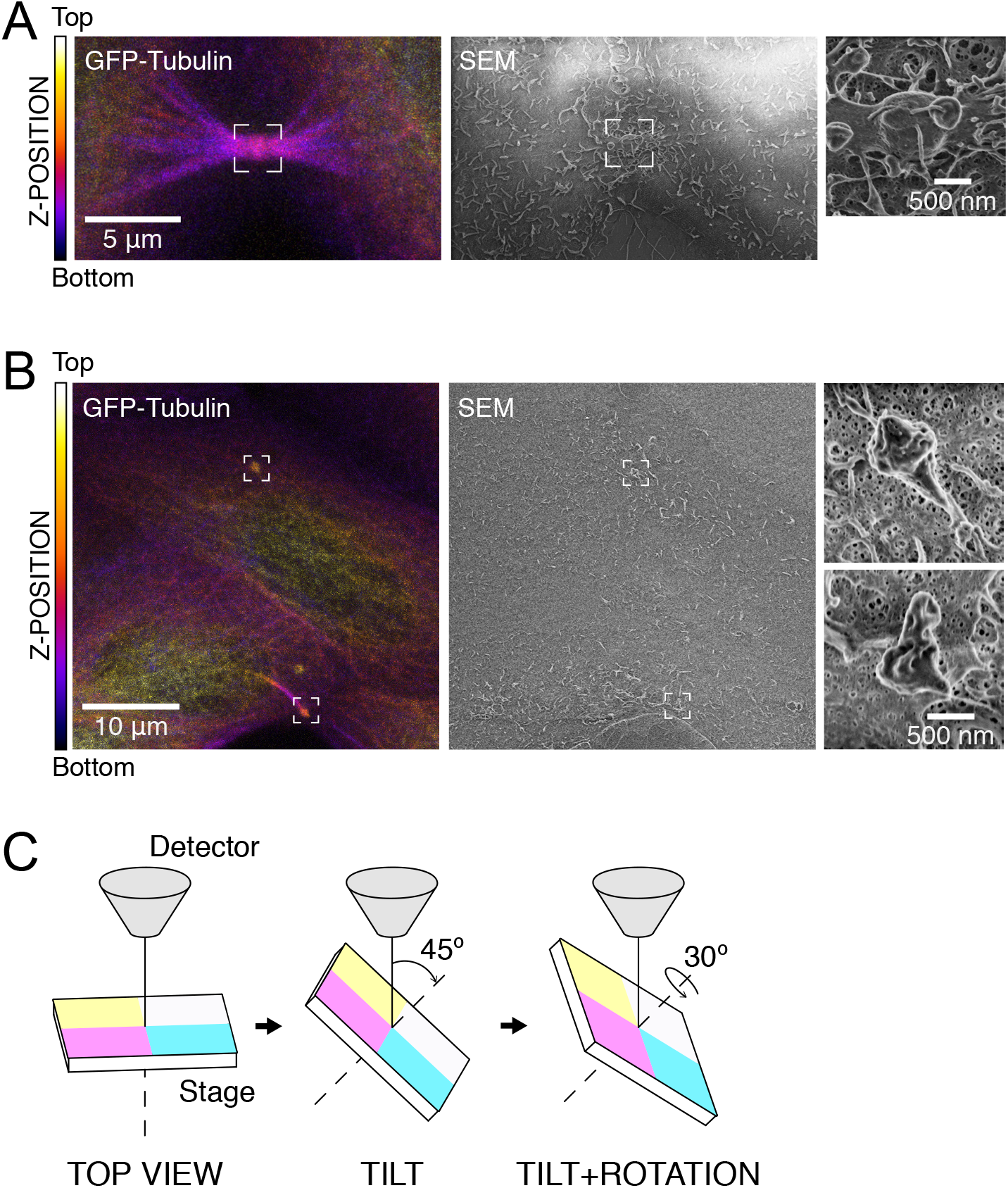
Procedure for correlative light and scanning electron microscopy. (**A**, **B**) Examples of CLEM imaging. Image of intercellular bridges during cytokinesis (A) and after abscission (B). In (A) there are microtubule bundles on both sides of the FB. The top structure in (B) corresponds to an MBR because microtubule bundles do not flank the FB; the structure at the bottom corresponds to a cleaved bridge that maintains microtubule bundles at the uncleaved arm. Confocal depth-coded color images of GFP-tubulin distribution (left), the corresponding SEM images (center), and enlargement of the boxed regions containing the structure of interest (right). The color scale used is indicated. (**C**) Procedure of side view image acquisition by SEM using tilting and rotation of the sample stage.

**Figure S3.**
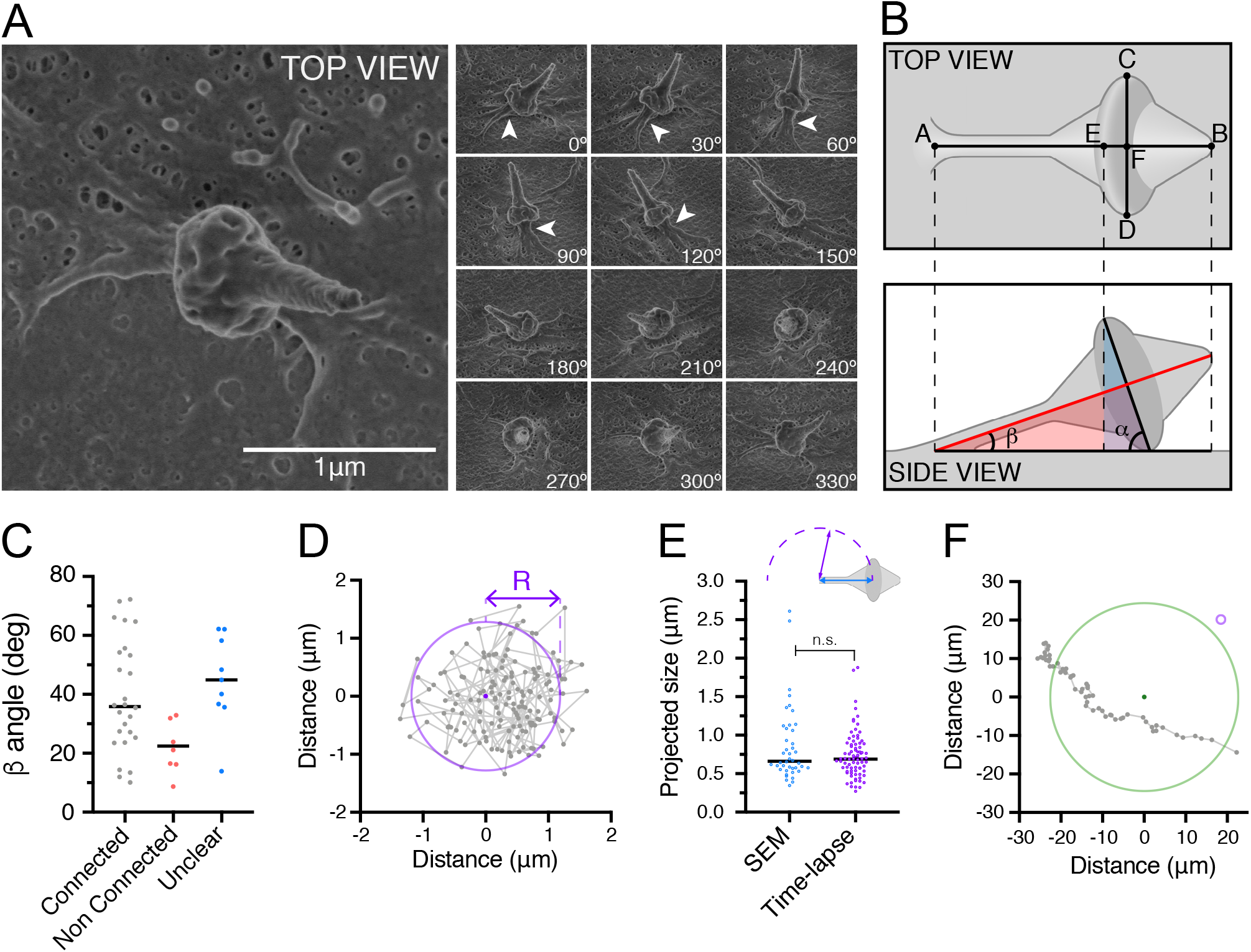
Tilting and rotation can reveal the existence of a connection. (**A**) Top-view image of an example of an MBR classified as an unclear case (left). The side-view series reveals that the MBR is connected to the plasma membrane (arrowhead; right panels). (**B**) Schematic illustrating how MBRs appear in top- and side-view images. MBR width was defined as the major axis of the FB (C-D line). The projected length of the long axis of the MBR is the distance between the two ends of the structure (A-B line). The intersection of the two lines defines the center of the FB (point F), which allows the measurement of the projected distance (A-F line) between the connection point and the center of the FB. The inclination angle (β) of the MBR with respect to the cell surface was derived from the inclination angle of the FB large axis (α), which was calculated from the distance between the FB rim (point E) and its center (point F). (**C**) β angle values for MBRs classified as connected, non-connected and unclear. (**D**) Example of the trajectory followed by an MBR. The purple dot represents the center of the trajectory. The circumference includes 95% of the dots. (**E**) Measurements of the projected distance between the connection point and the FB by SEM (n=42) compared with that calculated by the analysis of MBR trajectories obtained from time-lapse experiments (n=81). (**F**) Trajectory followed by a released MBR. The purple circle was drawn to be the same size as that in (D). Black bars represent median values. Probabilities are those associated with Mann-Whitney test.

**Figure S4.**
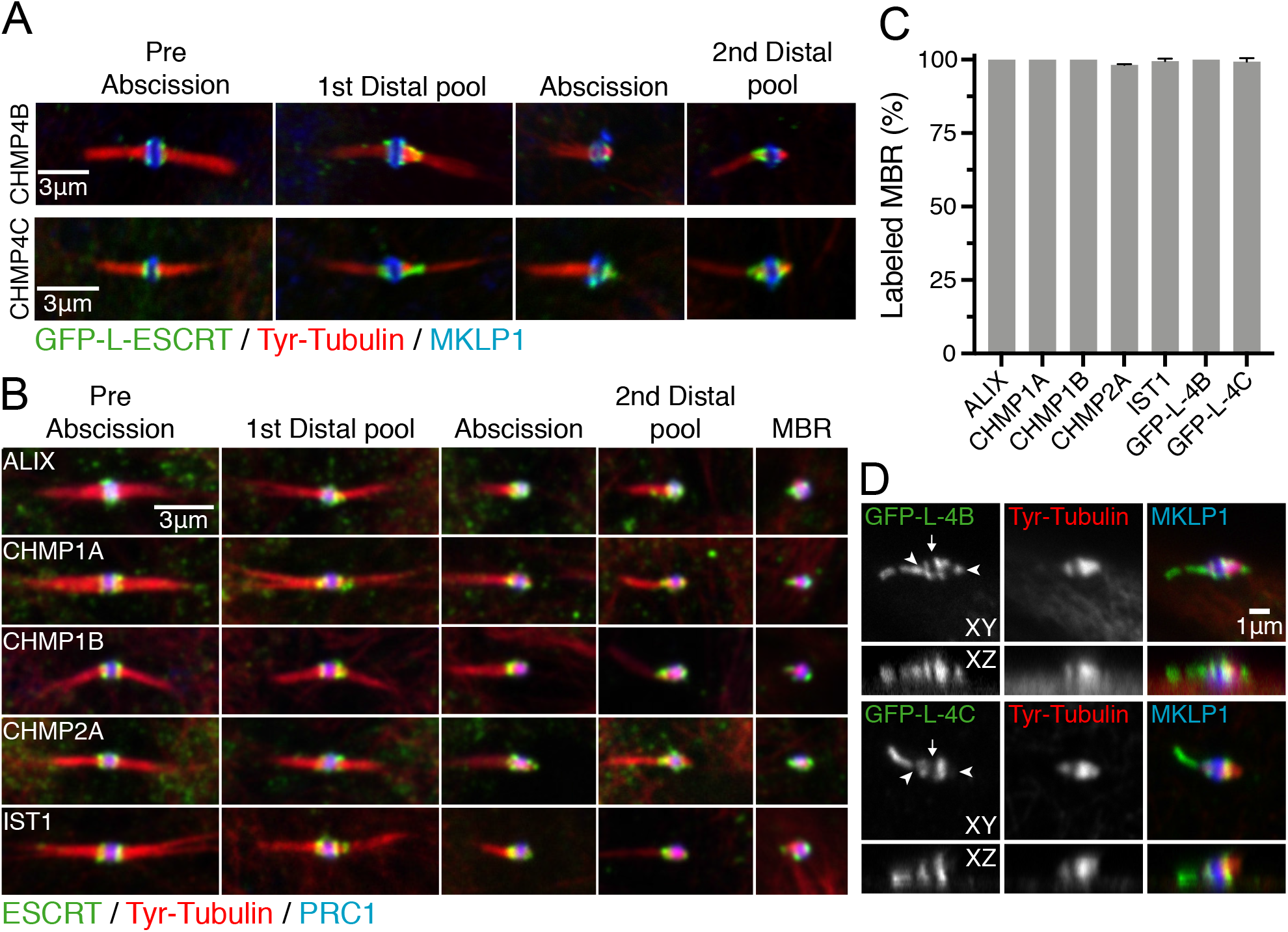
Distribution of ESCRT proteins at the MB and at the connection of the MBR with the plasma membrane. (**A**, **B**) Localization of GFP-L-CHMP4B and GFP-L-CHMP4C (A) and a panel of endogenous ESCRT proteins (B) at different stages of cytokinesis. (**C**) Percentage of MBRs positive for the indicated ESCRT proteins. The histogram represents the mean ± SD from three independent experiments (n=35-139). (**D**) Examples of MBRs showing uneven ESCRT distribution with an elongated pool of GFP-L-CHMP4B (top) and GFP-L-CHMP4C (bottom) in XY and XZ views. The arrow and the arrowheads indicate the FB and the MBR tips, respectively.

**Figure S5.**
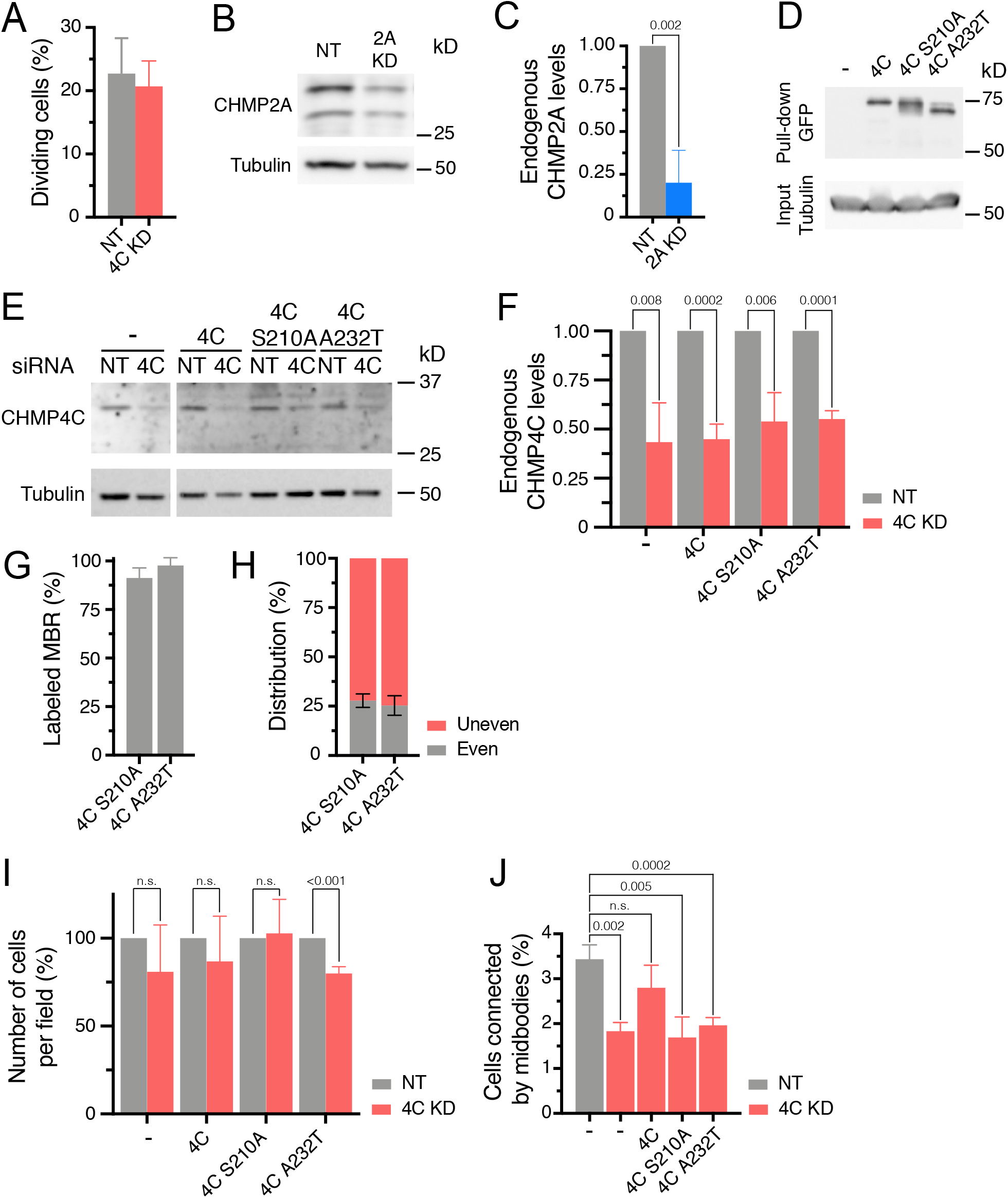
Effect of CHMP4C KD on the percentages of dividing cells and of cells connected by a MB. (**A**) Effect of CHMP4C KD on the frequency of dividing cells as determined by time-lapse experiments. The histogram represents the percentage of dividing cells relative to the initial number of cells. Three independent experiments (n=27-159 in control cells; n=8-95 in CHMP4C KD cells) were performed. (**B**) Representative immunoblot showing the effect of siCHMP2A on endogenous CHMP2A levels for the experiment shown in Figure 6D. (**C**) Quantification of CHMP2A KD by siCHMP2A. The histogram represents the levels of CHMP2A in siCHMP2A-transfected cells relative to that of cells transfected with control siNT. (**D**) Immunoblot of a GFP-trap experiment showing the relative expression levels of the indicated GFP-fused CHMP4C proteins. (**E**, **F**) Representative immunoblot (E) and quantification of endogenous CHMP4C levels (F) of siCHMP4-transfected cells expressing the indicated CHMP4C exogenous proteins. (**G**, **H**) Percentage of MBRs positive for the indicated CHMP4C mutants (n=58-80) (G). (H) Even or uneven distribution of the CHMP4C mutants in the MBR (n=37-56). (**I**) The percentage of CHMP4C KD cells (red bars) expressing the indicated exogenous CHMP4C proteins was calculated relative to their corresponding control cells (siNT, gray bars). (**J**) Percentage of cells connected by an MB in control cells and CHMP4C KD cells expressing the indicated exogenous CHMP4C proteins. The histograms in (A, C, F-J) show the mean ± SD from three independent experiments. Probabilities are those associated with unpaired two-tailed Student’s t-tests.

## Video captions

Video 1. **Side views of MBRs by SEM**. Complete rotation of a connected (left) and non-connected MBR (right). Numbers indicate the angle of rotation of the sample stage. This video corresponds to Figures 2A and B.

Video 2. **MBR movement on the apical surface**. 3D analysis of the movement of an MBR in a live cell expressing GFP-tubulin. This video corresponds to Figures 2D and E.

Video 3. **Dynamics of GFP-L-CHMP4B during and after abscission**. GFP-L-CHMP4B and Cherry-tubulin was tracked in cells during cytokinesis. This video is associated with Figure 4A.

Video 4. **CLEM images showing the presence of GFP-L-CHMP4C in the connection of the MBR with the plasma membrane**. Side view SEM images (left) and matching confocal images obtained by 3D reconstruction (right). The angle of rotation of the sample stage is indicated. This video corresponds to Figure 5B.

Video 5. **CLEM images showing the presence of GFP-L-CHMP4B in the connection of the MBR with the plasma membrane**. Side view SEM images (left) and matching confocal images obtained by 3D reconstruction (right). The angle of rotation of the sample stage is indicated. This video corresponds to Figure 5D.

Video 6. **GFP-L-CHMP4C and Cherry-tubulin distribution in a moving MBR**. (Left) GFP-L-CHMP4C and Cherry-tubulin fluorescence. (Right) The GFP-L-CHMP4C signal was pseudocolored using the indicated depth-color scale. This video corresponds to Figure 5F.

Video 7. **CHMP4C KD produces shedding of the MBR**. Control and CHMP4C KD cells expressing GFP-tubulin were recorded before and after abscission. This video is associated with Figure 6D.

